# Almond rhizosphere viral, prokaryotic, and fungal communities differed significantly among four California orchards and in comparison to bulk soil communities

**DOI:** 10.1101/2023.06.03.543555

**Authors:** Anneliek M. ter Horst, Temiloluwa V. Adebiyi, Daisy A. Hernandez, Jane D. Fudyma, Joanne B. Emerson

## Abstract

Characterization of rhizosphere microbiomes and their interactions is essential to a holistic understanding of plant health in support of sustainable agriculture. Viruses are a key, understudied component of rhizosphere microbiomes, with potential impacts on both plant-beneficial and -pathogenic organisms through infection. In this study, we sampled rhizospheres and bulk soils associated with 15 almond trees in four California orchards and generated viromic, 16S rRNA gene, and ITS1 amplicon sequencing datasets to compare viral, prokaryotic, and fungal communities. In total, 10,440 viral operational taxonomic units (vOTUs), 16,146 bacterial and archaeal OTUs, and 6,684 fungal OTUs were recovered. All three community types differed most significantly among the four orchards and secondarily between bulk and rhizosphere soils. Despite compositional differences, no significant differences in richness were observed between bulk and rhizosphere soils for any of the studied biota. Overall, viruses, prokaryotes, and fungi shared similar beta-diversity patterns in almond rhizospheres and bulk soils on a regional scale, counter to recently observed decoupling between viral and prokaryotic community biogeographic patterns in a variety of bulk soils.

## Introduction

Almonds are a major cash crop in the United States, particularly in California, which produces more than 83% of the world’s almonds^1^. Given the worldwide importance of the specialty crop, it is necessary to minimize losses due to disease through the understanding of the microbial drivers behind the development of disease in perennial crops. Knowledge of almond-associated bacterial, fungal, and viral communities and their interactions will facilitate a more holistic understanding of almond trees in healthy and diseased states. Rhizosphere root zones, in particular, are important gateways for plant-microbe interactions, as plants secrete nutrients and carbon into the rhizosphere, attracting plant-beneficial and other microbes ^2–6^. Beneficial symbiotic relationships among plants, fungi, and bacteria in the rhizosphere have been extensively studied, but very little is known about rhizosphere viral communities^7^.

Recent viral community ecological studies in bulk soil offer insight on biogeographical patterns that we might expect to also occur in the rhizosphere. For example, oil viral communities are highly diverse^8,9^, differ over short spatial distances both within the same field^10^ and at regional scales^11,12^, and they share some similarity by habitat^13^. Early insights into rhizosphere viral community composition suggest some ecological patterns similar to their host bacteria, fungi, and/or other organisms, such as significant differences between bulk and rhizosphere soil compartments. For example, a comparison of four bulk and four rhizosphere soil samples showed significant differences in viral communities between these compartments in a maize system^14^, and a recent study of viral communities of the oilseed rape rhizosphere found that crop rotation practice sometimes had a greater effect on viral community composition than soil compartment^15^. Significant differences in RNA viral communities across rhizosphere, bulk soil, and detritusphere (litter-associated) compartments were also observed in greenhouse experiments on wild oat grasses^16^. Given the stark differences in viral and bacterial community compositional patterns previously established in a variety of soils^10,17^, as well as the known soil viral community heterogeneity at local and regional scales^11,12,17,18^, it is unknown whether microbial community rhizosphere beta-diversity patterns reflect those of their viruses as a general rule.

Here we sampled a total of 15 almond tree rhizospheres and their accompanying bulk soils from four orchards in California (Figure 1A). From these 30 samples, we generated viral size-fraction metagenomic (viromic), as well as 16S rRNA gene and ITS1 amplicon datasets to compare viral, bacterial, and fungal beta-diversity patterns in bulk and rhizosphere soils within and among orchards on a regional scale.

**Figure 1:**
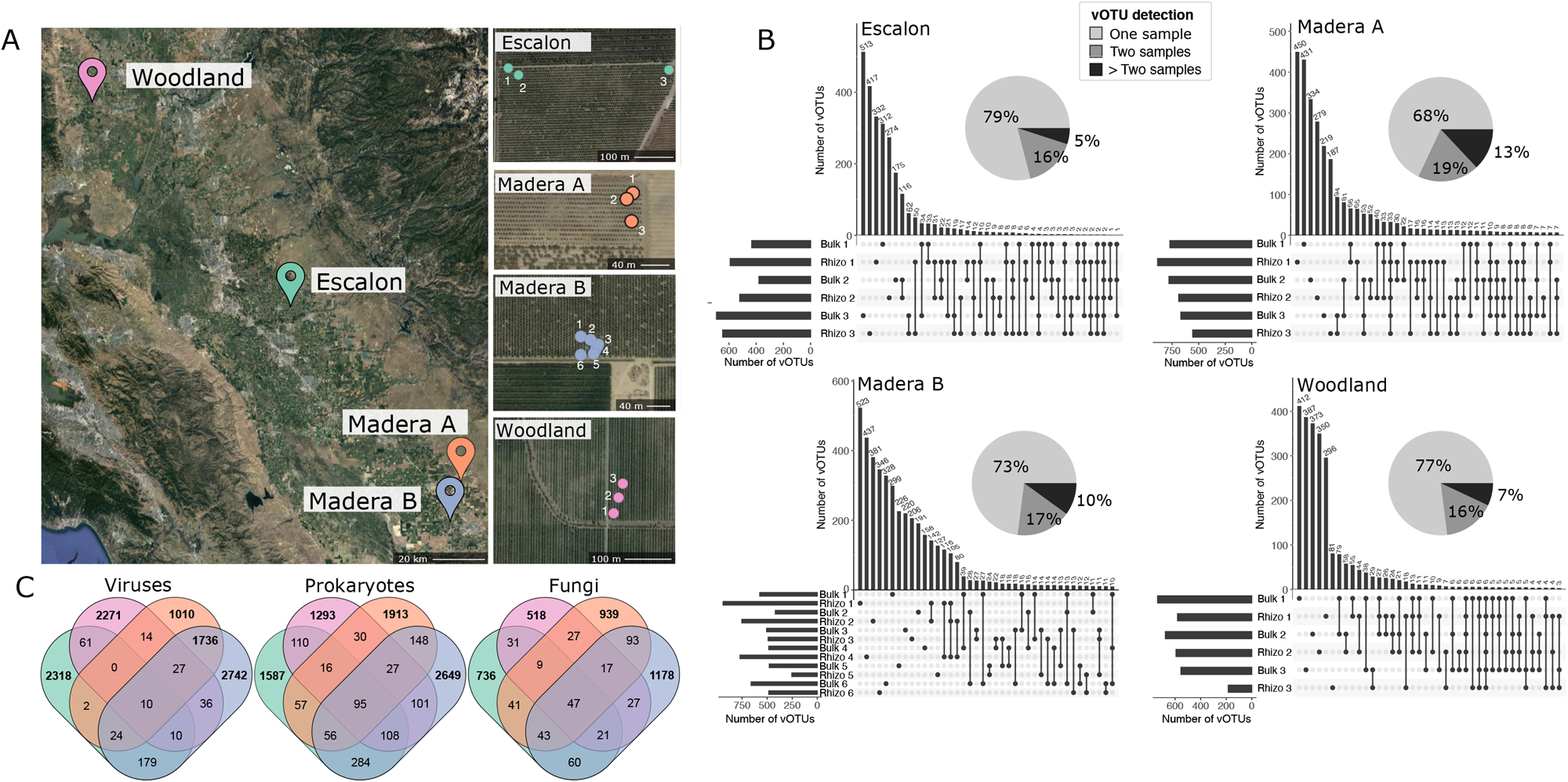
Sampling locations and detection patterns for viral, prokaryotic, and fungal species in California almond orchards. **A)** Map of the locations of the four orchards (left) and locations of each pair of samples (tree roots and accompanying bulk soil) within each orchard (right). **B)** UpSet plot indicating vOTU detection patterns and pie chart indicating how many of the vOTUs were present in one, two, or more than two samples for each of the four orchards. **C)** Venn diagrams of the numbers of vOTUs (Viruses), microbial 16S rRNA gene OTUs (Prokaryotes), and fungal ITS amplicon OTUs (Fungi) detected within or across the four orchards. Numbers >1,000 (viruses and prokaryotes) or >500 (fungi) are in bold for ease of interpretation.

## Results and Discussion

The four sampled orchards were located in three areas spanning an approximately 1,585 km^2^ region, with two of the orchards (Madera A and B) in the same county 16 km apart (Figure 1A). Rhizospheres from three healthy trees and their nearby bulk soils were sampled from each orchard, and three additional diseased tree rhizospheres and associated bulk soils were sampled from Madera B, for a total of 30 samples (15 bulk soil, 15 rhizosphere, Figure 1A). From each sample, we generated viral-size fraction metagenomes (viromes) to study viral communities, as well as 16S rRNA gene and ITS1 amplicon sequencing datasets to study bacterial and fungal communities (Supplementary Figure 1), respectively. We used the viromes to identify biogeographical patterns in viral community composition and all datasets to compare viral, bacterial, and fungal beta-diversity patterns.

From the 30 viromes, 46,501 viral contigs were assembled and clustered into 41,305 viral operational taxonomic units (vOTUs, ≥ 10 kbp, ≥ 95% average nucleotide identity, representing approximately species-level taxonomy^19^). Of these vOTUs, 10,440 were classified as medium- or high-quality by VIBRANT^20^ and used for further analysis. Many vOTUs (62%) were only recovered in a single virome in the dataset, similar to a previous regional-scale comparison of viral communities in four natural soil habitats, in which 81% of vOTUs were only found in one of 30 viromes^11^. Within each orchard, the vast majority of vOTUs (74% on average) was only detected in a single virome, indicating substantial viral community heterogeneity within the same field (Figure 1B). Co-occurring vOTUs in the same orchard were most often found either in paired rhizosphere-bulk soil samples from the same tree (12.7%) or in the same habitat type (bulk-bulk (12.8%) or rhizosphere-rhizosphere (17.9%) pairs), indicating that both proximity and habitat were strong contributors to viral community composition in almond orchards. Of the 2,099 vOTUs shared between two or more orchards, the majority (86%) were shared between the two Madera orchards, which were in closest proximity, presumably reflecting the greater dispersal potential between and/or more similar environmental conditions at these two sites.

Although the fraction of vOTUs shared between the two Madera orchards was higher than for bacterial OTUs or fungal OTUs, overall, similar patterns were revealed for 16S rRNA gene amplicon sequence variants (OTUs) and fungal operational taxonomic units (OTUs), of which only 8% and 10%, respectively, were detected in more than one orchard (Figure 1C).

Viral, prokaryotic, and fungal communities each differed primarily among the four orchards (meaning, by location, Figure 2A-C). Secondarily, communities of all three types were significantly different in bulk compared to rhizosphere soils (Figure 2D-F), as previously shown many times for bacteria and fungi (e.g., ^4^). Despite differences in community composition, richness was not significantly different between bulk and rhizosphere soils for any of these biota (Supplementary Figure 2, Student’s T-tests, p > 0.01). Viral community beta-diversity patterns were significantly correlated with those of both bacteria and fungi (Mantel tests, p < 0.01), indicating that all three groups may be responding to similar environmental cues and/or to each other. Together, these results suggest that viruses, prokaryotes, and fungi shared more similar beta-diversity patterns in rhizosphere and bulk soils than might have been predicted from prior studies of viral and prokaryotic communities in bulk soils alone^10,17^. As expected from prior regional studies in bulk soils^11^, rhizosphere viral communities were significantly different among orchards on a regional scale. However, prokaryotic and fungal communities also shared these patterns, suggesting the potential for greater coupling of viral and host beta-diversity patterns in rhizospheres, compared to the decoupled patterns recently observed in bulk soils. Given the substantial differences in community composition between orchards and the tendency for many populations (from all three types of biota) to be recovered in single orchards or even single samples here, future studies of rhizosphere viral community ecology would benefit from increased spatiotemporal resolution, for example, longitudinal studies of multiple plants in the same field and/or in greenhouse experiments.

**Figure 2:**
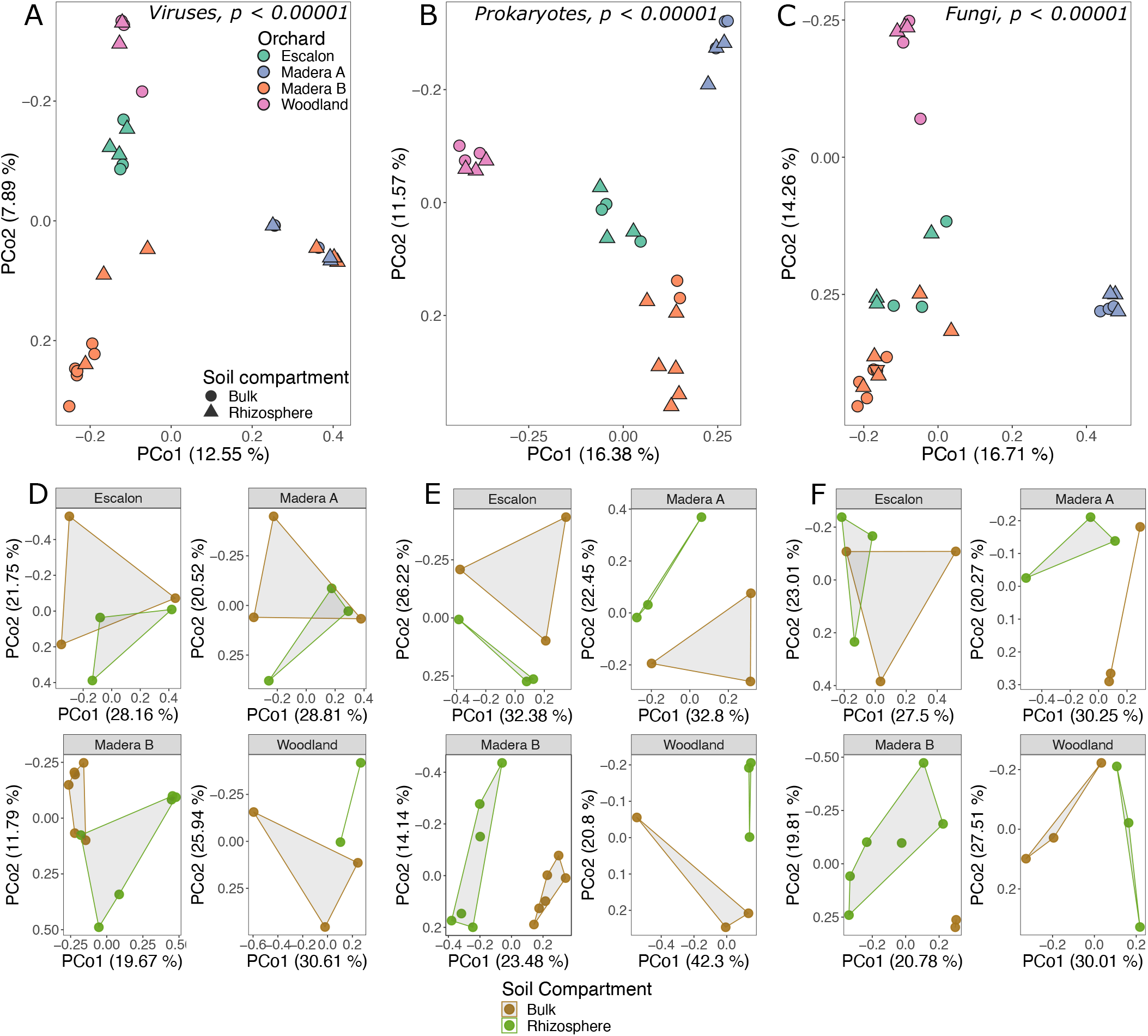
Community compositional structuring of viral, prokaryotic, and fungal communities across and within orchards. **A, B, C:** Principal coordinates analyses performed on Bray-Curtis dissimilarities of **A)** vOTUs, **B)** 16S rRNA gene OTUs, and **C)** ITS OTUs, colored by the orchard from which the sample was taken. **D, E, F:** Principal coordinates analyses performed on Bray-Curtis dissimilarities of **D)** vOTUs, **E)** 16S rRNA gene OTUs and **F)** ITS OTUs, in separate plots by orchard and colored by soil compartment (bulk soil or rhizosphere soil). For all plots, percentages along the axes indicate the percent variance explained.

## Methods

### Sample collection

Collection of bulk soil and rhizosphere samples occurred at four almond orchards in California: Escalon (N37.48.754, W120.56.187), Madera A (N37.07.915, W120.06.953), Madera B (N36.59.577, W120.10.185), and Woodland (N38.35.184, W121.53.337) (Supplementary table 1). One orchard was sampled in Escalon, one in Woodland, and two in Madera, with the Madera orchards 16 km apart and all orchards spread across a 1,585 km^2^ area. Rhizosphere and nearby bulk soil samples were collected from three healthy trees per orchard and also from three diseased trees infected with Ceratocystis cankers at the Madera B orchard, for a total of 30 samples. Rhizosphere soils were collected from three roots per tree. Roots were collected by excavating near-surface root systems and cutting three roots out of the soil, using a knife sterilized first with 70% ethanol and second with 10% bleach. Rhizosphere soils were defined as root-attached soil remaining after shaking the roots vigorously. Bulk soil was defined as soil 2 m from the trunk of an adjacent tree with a rhizosphere sample, and a single core 7 cm in diameter and 15 cm deep was collected for each bulk soil sample. Bulk soil and root samples were transported to the lab on ice and stored at 4 °C until processing. Rhizosphere soil separation and viromic and total DNA extraction from bulk and rhizosphere soils was completed for all samples within 48 hours of collection.

### Viromic DNA extraction and sequencing

Rhizosphere viral purification was performed by washing the roots in a 50 mL conical tube with 30 mL of protein-amended phosphate buffered saline (PPBS) buffer ^21^. Bulk soil was put through an 8 mm sieve to remove root material and then placed in a 50 mL tube with 30 mL of PPBS buffer. From this point, bulk and rhizosphere viromic samples were treated the same way. Tubes were shaken for 30 minutes at 300 rpm at 4 °C, then spun for 10 minutes at 4,000 x g. The supernatant was retained and centrifuged at 10,000 x g for 8 minutes. The supernatant was then passed through a 0.22 μm filter to remove most cellular components and larger soil particles, and the filtered supernatant was ultracentrifuged (Beckman LE-80K) at 35,000 rpm for 2.25 hours. After the supernatant was decanted, each pellet was solubilized in 100 μL of sterile water and then treated with 10 μL DNase (Promega, Madison, WI, USA) for 30 minutes at 37 °C. 10 μL DNase stop reagent (Promega) was added to stop the reaction. DNA extraction from the viral fraction was performed with the PowerSoil Pro Kit (QIAGEN, Hilden, Germany), following the manufacturer’s instructions, with an added step of a 10-minute incubation at 65 ºC before the bead-beating step. The extracted DNA underwent KAPA DNA Hyper library construction and was sequenced at the UC Davis DNA Technologies Core on the Illumina NovaSeq6000 platform (150 cycles paired end). All viromes were sequenced to an approximate depth of 10 Gbp.

### Total DNA extraction and amplicon sequencing

Total DNA was extracted from 0.25 g bulk or rhizosphere soil per sample with the PowerSoil Pro Kit (QIAGEN, Hilden, Germany), using the same lysis and extraction protocol as for the viromes. For rhizosphere soil, soil that was adhered to the roots was brushed off and placed in the extraction tube. For each sample, amplicons were generated for 16S rRNA gene and ITS1 amplicon sequencing via PCR reactions with the extracted DNA as a template.

Construction of 16S rRNA gene amplicon libraries followed a previously described dual-indexing strategy ^22,23^. To target the V4 region of the 16S rRNA gene, universal primers 515F and 806R were used, using the following PCR protocol: an initial denaturation step at 98 ºC for 2 min, followed by 30 cycles of 98 ºC for 20 s, 50 ºC for 30 s and 72 ºC for 45 s, and a final extension step at 72 ºC for 10 min.

To amplify the ITS1 region, we used the universal primers ITS1-F and ITS2 ^24–26^ and the following PCR program: an initial denaturation step at 95 °C for 2 min, followed by 35 cycles of 95 °C for 20 s, 50 °C for 30 s, and 72 °C for 50 s, followed by a final extension at 72 °C for 10 min. All PCR reactions were performed using the Platinum Hot Start PCR Master Mix (Invitrogen). Libraries were cleaned using AmpureXP magnetic beads (Beckman Coulter), quantified (Qubit 4 fluorometer), and pooled in equimolar concentrations. Paired-end sequencing (250 bp) was performed on the MiSeq platform (Illumina) at the UC Davis DNA Technologies Core.

### Bioinformatic processing of viromes

Trimmomatic ^27^ was used to quality-filter the reads, and adapters and PhiX were removed. MEGAHIT ^28^ was used to assemble reads, using default parameters. Dereplication was performed with dRep, at 95% ANI with a minimum coverage threshold of 85%, using the ANImf algorithm ^29^. VIBRANT was used to predict viral contigs, in virome mode ^20^. Bowtie2 ^30^ was used to map reads to vOTU sequences, using the sensitive mode. CoverM (https://github.com/wwood/CoverM^31^ was used to create a coverage table, using default parameters.

### Bioinformatic processing of amplicon sequencing datasets

Paired-end read assembly into single sequences was done using PANDAseq v2.9 ^32^, and chimeric sequence removal was done using usearch v6.1 ^33^. OTU clustering at 97% sequence identity was done using the QIIME ^34^ implementation of UCLUST v1.2.22 ^33^, against the SILVA database v132 ^35^ for 16S rRNA gene sequences and against the UNITE database v2021-05-10 for ITS1 sequences ^36^.

### Statistical analyses and figure generation

All statistical analyses were performed using R v4.1.0 ^37^, using the mean coverage vOTU abundance table for viruses and the OTU abundance tables for prokaryotes and fungi. Bray-Curtis dissimilarities were calculated on log-transformed relative abundances, using the vegdist function from Vegan v2.6-2 ^38^, and principal coordinates analyses were performed with the pcoa() function from ape v5.4-2 ^39^. All maps were generated using Google Earth, and all other plots were created using the R package ggplot2 v3.3.5 ^40^.

## Data availability

Raw sequencing reads for viromes, 16S rRNA and ITS rRNA are available on NCBI under bioproject number PRJNA915061 and vOTU sequences are available on Dryad within the PIGEONv2.0 database (https://doi.org/10.25338/B8C934).100 m

## References

1. Marvinney, E., Kendall, A., Brodt, S. & Others. A comparative assessment of greenhouse gas emissions in California almond, pistachio, and walnut production. in Proceedings of the 9th International Conference on Life Cycle Assessment in the Agri-Food Sector 761–771 (2014).

2. Ling, N., Wang, T. & Kuzyakov, Y. Rhizosphere bacteriome structure and functions. Nat. Commun. 13, 836 (2022).

3. Bakker, P. A. H. M., Pieterse, C. M. J., de Jonge, R. & Berendsen, R. L. The Soil-Borne Legacy. Cell 172, 1178–1180 (2018).

4. Nuccio, E. E. et al. Niche differentiation is spatially and temporally regulated in the rhizosphere. ISME J. 14, 999–1014 (2020).

5. Philippot, L., Raaijmakers, J. M., Lemanceau, P. & van der Putten, W. H. Going back to the roots: the microbial ecology of the rhizosphere. Nat. Rev. Microbiol. 11, 789–799 (2013).

6. Mendes, R., Garbeva, P. & Raaijmakers, J. M. The rhizosphere microbiome: significance of plant beneficial, plant pathogenic, and human pathogenic microorganisms. FEMS Microbiol. Rev. 37, 634–663 (2013).

7. Pratama, A. A., Terpstra, J., de Oliveria, A. L. M. & Salles, J. F. The Role of Rhizosphere Bacteriophages in Plant Health. Trends Microbiol. 28, 709–718 (2020).

8. Roux, S. & Emerson, J. B. Diversity in the soil virosphere: to infinity and beyond? Trends Microbiol. 30, 1025–1035 (2022).

9. Williamson, K. E., Fuhrmann, J. J., Wommack, K. E. & Radosevich, M. Viruses in Soil Ecosystems: An Unknown Quantity Within an Unexplored Territory. Annu Rev Virol 4, 201–219 (2017).

10. Santos-Medellín, C. et al. Spatial turnover of soil viral populations and genotypes overlain by cohesive responses to moisture in grasslands. Proc. Natl. Acad. Sci. U. S. A. 119, e2209132119 (2022).

11. Durham, D. M. et al. Substantial differences in soil viral community composition within and among four Northern California habitats. ISME Communications 2, 1–5 (2022).

12. Hillary, L. S., Adriaenssens, E. M., Jones, D. L. & McDonald, J. E. RNA-viromics reveals diverse communities of soil RNA viruses with the potential to affect grassland ecosystems across multiple trophic levels. ISME Commun 2, 34 (2022).

13. Ter Horst, A. M. et al. Minnesota peat viromes reveal terrestrial and aquatic niche partitioning for local and global viral populations. Microbiome 9, 233 (2021).

14. Bi, L. et al. Diversity and potential biogeochemical impacts of viruses in bulk and rhizosphere soils. Environ. Microbiol. 23, 588–599 (2021).

15. Muscatt, G. et al. Crop management shapes the diversity and activity of DNA and RNA viruses in the rhizosphere. Microbiome 10, 181 (2022).

16. Starr, E. P., Nuccio, E. E., Pett-Ridge, J., Banfield, J. F. & Firestone, M. K. Metatranscriptomic reconstruction reveals RNA viruses with the potential to shape carbon cycling in soil. Proc. Natl. Acad. Sci. U. S. A. 116, 25900–25908 (2019).

17. Santos-Medellin, C. et al. Viromes outperform total metagenomes in revealing the spatiotemporal patterns of agricultural soil viral communities. ISME J. 15, 1956–1970 (2021).

18. Ter Horst, A. M., Fudyma, J. D., Sones, J. L. & Emerson, J. B. Dispersal, habitat filtering, and eco-evolutionary dynamics as drivers of local and global wetland viral biogeography. bioRxiv 2023.04.28.538735 (2023) doi:10.1101/2023.04.28.538735.

19. Roux, S. et al. Minimum Information about an Uncultivated Virus Genome (MIUViG). Nat. Biotechnol. 37, 29–37 (2019).

20. Kieft, K., Zhou, Z. & Anantharaman, K. VIBRANT: automated recovery, annotation and curation of microbial viruses, and evaluation of viral community function from genomic sequences. Microbiome 8, 90 (2020).

21. Göller, P. C., Haro-Moreno, J. M., Rodriguez-Valera, F., Loessner, M. J. & Gómez-Sanz, E. Uncovering a hidden diversity: optimized protocols for the extraction of dsDNA bacteriophages from soil. Microbiome vol. 8 Preprint at https://doi.org/10.1186/s40168-020-0795-2 (2020).

22. Caporaso, J. G. et al. QIIME allows analysis of high-throughput community sequencing data. Nat. Methods 7, 335–336 (2010).

23. Edwards, J., Santos-Medellín, C. & Sundaresan, V. Extraction and 16S rRNA Sequence Analysis of Microbiomes Associated with Rice Roots. Bio Protoc 8, e2884 (2018).

24. Gardes, M. & Bruns, T. D. ITS primers with enhanced specificity for basidiomycetes--application to the identification of mycorrhizae and rusts. Mol. Ecol. 2, 113–118 (1993).

25. White, T. J., Bruns, T., Lee, S., Taylor, J. & Others. Amplification and direct sequencing of fungal ribosomal RNA genes for phylogenetics. PCR protocols: a guide to methods and applications 18, 315–322 (1990).

26. Agler, M. T. et al. Microbial Hub Taxa Link Host and Abiotic Factors to Plant Microbiome Variation. PLoS Biol. 14, e1002352 (2016).

27. Bolger, A. M., Lohse, M. & Usadel, B. Trimmomatic: a flexible trimmer for Illumina sequence data. Bioinformatics 30, 2114–2120 (2014).

28. Li, D., Liu, C.-M., Luo, R., Sadakane, K. & Lam, T.-W. MEGAHIT: an ultra-fast single-node solution for large and complex metagenomics assembly via succinct de Bruijn graph. Bioinformatics 31, 1674–1676 (2015).

29. Olm, M. R., Brown, C. T., Brooks, B. & Banfield, J. F. dRep: a tool for fast and accurate genomic comparisons that enables improved genome recovery from metagenomes through de-replication. ISME J. 11, 2864–2868 (2017).

30. Longmead, B. & Salzberg, S. L. Fast gapped-read alignment with Bowtie2. Nat. Methods 9, 357–359 (2012).

31. Woodcroft, B. J. CoverM: Read coverage calculator for metagenomics. (Github).

32. Masella, A. P., Bartram, A. K., Truszkowski, J. M., Brown, D. G. & Neufeld, J. D. PANDAseq: paired-end assembler for illumina sequences. BMC Bioinformatics 13, 31 (2012).

33. Edgar, R. C. Search and clustering orders of magnitude faster than BLAST. Bioinformatics 26, 2460–2461 (2010).

34. Bolyen, E. et al. Reproducible, interactive, scalable and extensible microbiome data science using QIIME 2. Nat. Biotechnol. 37, 852–857 (2019).

35. Quast, C. et al. The SILVA ribosomal RNA gene database project: improved data processing and web-based tools. Nucleic Acids Res. 41, D590–6 (2013).

36. Abarenkov, K. et al. UNITE QIIME release for fungi. (2021) doi:10.15156/BIO/1264708.

37. RCore, T. R: A language and environment for statistical computing. R Foundation for Statistical Computing, Vienna, Austria. Preprint at (2016).

38. Oksanen, J. et al. vegan: Community Ecology Package. R package version 2.5-2. 2018. Preprint at (2018).

39. Paradis, E. & Schliep, K. ape 5.0: an environment for modern phylogenetics and evolutionary analyses in R. Bioinformatics 35, 526–528 (2019).

40. Wickham, H. ggplot2: elegant graphics for data analysis. data. (2016).

