## Supplementary Figure 1 for "Almond rhizosphere viral, prokaryotic, and fungal communities differed significantly among four California orchards and in comparison to bulk soil communities"

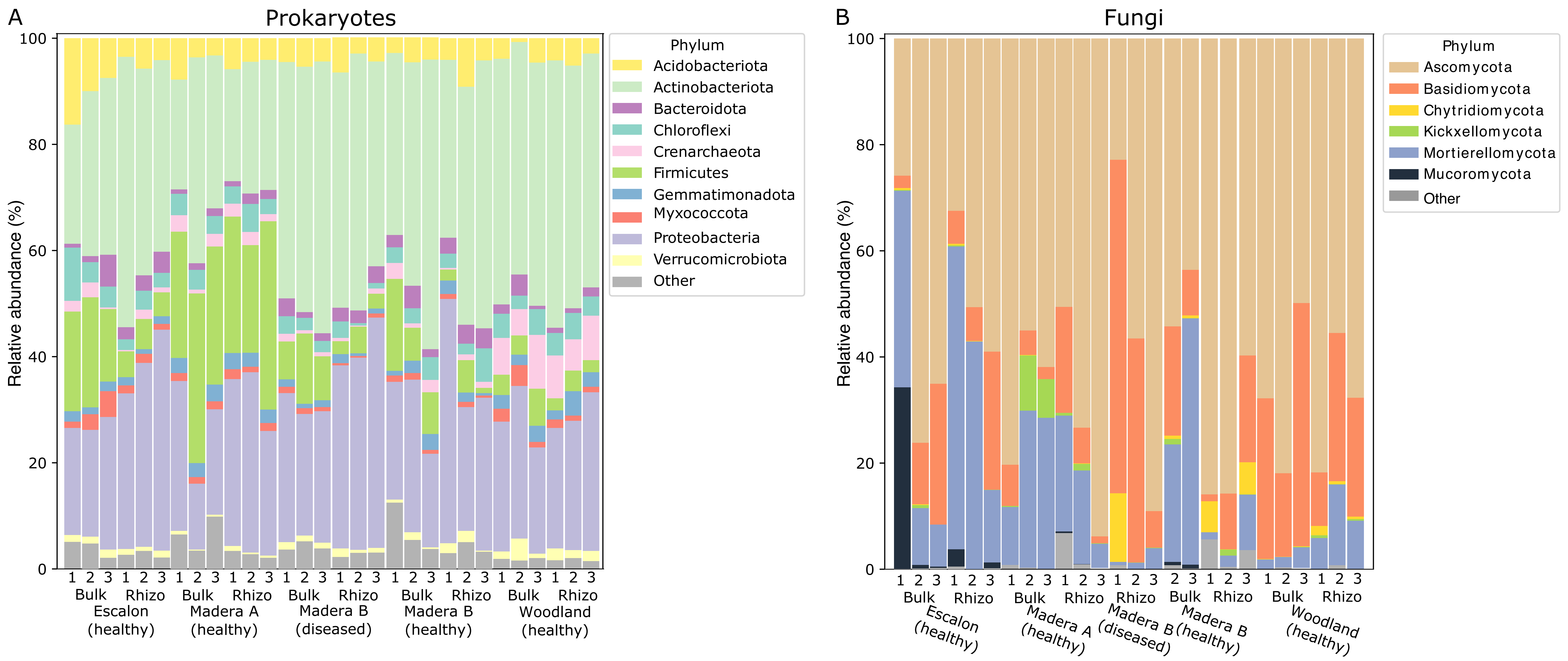


**Supplementary figure 1:** Relative abundances of prokaryotes and fungi in each sample. **A)** Relative abundances of 16S rRNA gene OTUs (summed at the phylum level) in each of the samples. **B)** Relative abundances of ITS OTUs (summed at the phylum level) in each of the samples.
