## Supplementary Figure 2 for "Almond rhizosphere viral, prokaryotic, and fungal communities differed significantly among four California orchards and in comparison to bulk soil communities"

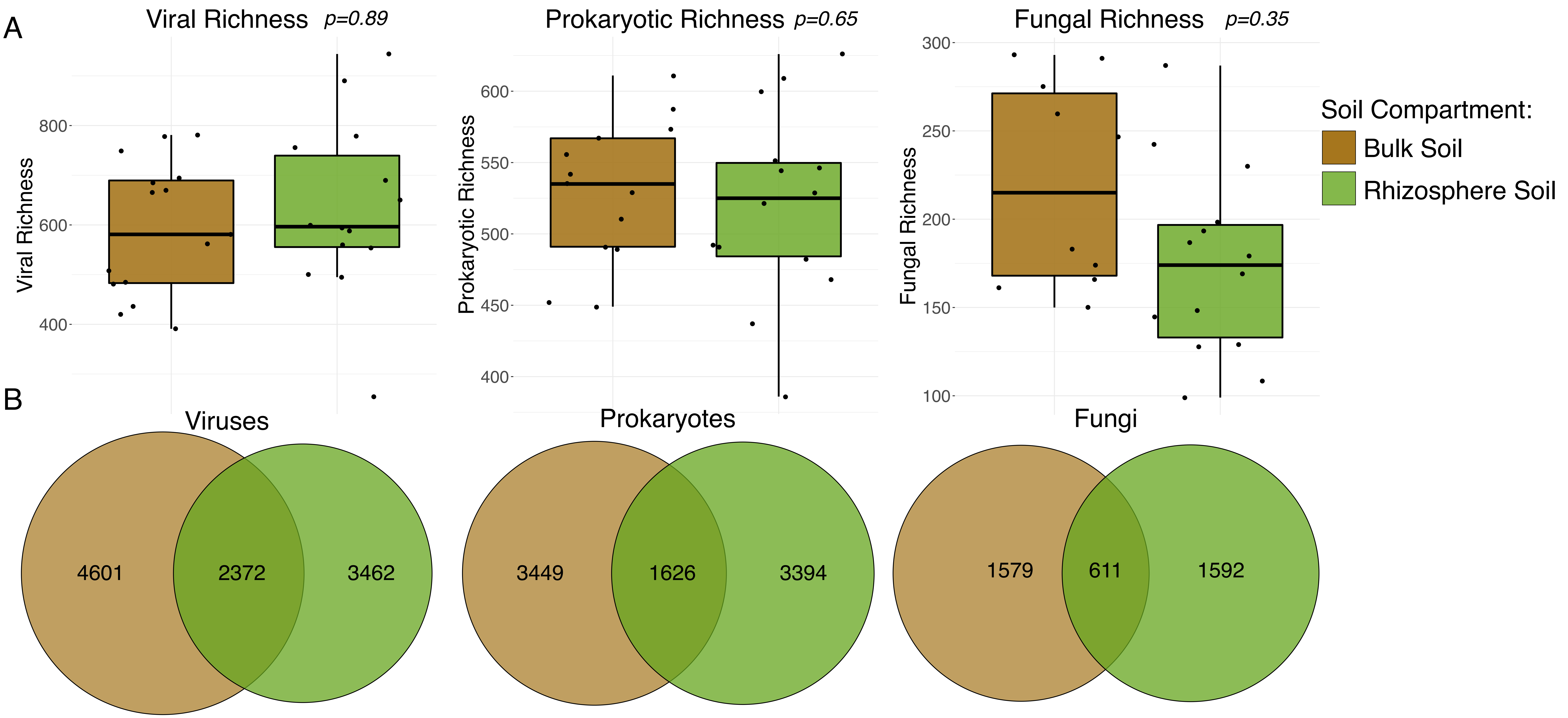


**Supplementary figure 2: A**) Richness (number of vOTUs or OTUs) for viral (vOTUs), prokaryotic (16S rRNA gene OTUs), and fungal (ITS OTUs) communities. Each point is one sample, lines are medians, boxes are the first interquartile range, and whiskers represent the 95th percentile. P-values are Student's T-test results, comparing richness values in bulk versus rhizosphere soils (significant when $p < 0.05$). **B)** Venn diagrams of the number of vOTUs (viruses) or OTUs (prokaryotes and fungi) detected in one or both soil compartments.
